# Whole genome sequencing based differentiation between re-infection and relapse in Indian patients with tuberculosis recurrence, with and without HIV co-infection

**DOI:** 10.1101/2020.07.29.227926

**Authors:** Sivakumar Shanmugam, Nathan L Bachmann, Elena Martinez, Ranjeeta Menon, Gopalan Narendran, Sujantha Narayanan, Srikanth Tripathy, K R Uma Devi, Shailendra Sawleshwarkar, Ben Marais, Vitali Sintchenko

## Abstract

Differentiation between relapse and reinfection in cases with tuberculosis (TB) recurrence has important implications for public health, especially in patients with human immunodeficiency virus (HIV) co-infection. Forty-one paired *M. tuberculosis* isolates collected from 20 HIV-positive and 21 HIV-negative patients, who experienced TB recurrence after previous successful treatment, were subjected to whole genome sequencing (WGS) in addition to spoligotyping and mycobacterial interspersed repeat unit (MIRU) typing. Comparison of *M. tuberculosis* genomes indicated that 95% of TB recurrences in the HIV-negative cohort were due to relapse, while the majority of TB recurrences (75%) in the HIV-positive cohort was due to re-infection (P=0.0001). Drug resistance conferring mutations were documented in four pairs (9%) of isolates associated with relapse. The high contribution of re-infection to TB among HIV patients warrants further study to explore risk factors for TB exposure in the community.

## Introduction

Recurrence of tuberculosis (TB) can result from reactivation of an infection that was only partially cleared with past treatment (endogenous relapse) or from exogenous re-infection with a new strain of *Mycobacterium tuberculosis*. Over the years several *M. tuberculosis* typing methods have been developed to assist differentiation between these two causes, such as restriction fragment length polymorphism (RFLP) typing, spoligotyping and mycobacterial interspersed repeat unit (MIRU) typing [1]. In recent times whole genome sequencing (WGS) has demonstrated superior resolution to identify TB transmission pathways [2] and to differentiate TB relapse from re-infection in low incidence settings [3, 4].

The human immunodeficiency virus (HIV) epidemic poses a significant obstacle to global TB control, mainly because the immunosuppression induced by HIV increases the risk of TB reactivation and disease progression after recent infection [5]. Globally, TB is the leading cause of death in HIV-infected patients and India is estimated to have the highest number of new TB cases in the world (>20% of all cases), as well as the highest number of new HIV infections in the Asia-Pacific Region [6]. TB recurrence is a major problem in high burden countries like India and the risk of TB recurrence is increased by HIV co-infection [7].

It has been observed that relapse is a more common cause of TB recurrence in HIV-negative compared to HIV-positive patients [8, 9]. Given that accurate differentiation between relapse and re-infection is important to consider optimal treatment and infection control measures, we aimed to critically examine the relative contribution of relapse and re-infection to TB recurrence in HIV-positive and -negative patients using WGS.

## Methods

### Study population

We performed retrospective sequencing of *M. tuberculosis* recovered from specimens collected during TB treatment trials conducted in Chennai, India. Specimens selected for this analysis were identical to those used in a previous ‘case-control’ study comparing TB relapse in HIV-positive and -negative patients using spoligo- and mycobacterial interspersed repeat unit (MIRU)–12 typing [9]; except for six instances where banked DNA samples failed quality checks. Participants were newly diagnosed pulmonary TB cases enrolled into prospective treatment trials. The first cohort consists of 21 HIV-negative patients with pulmonary tuberculosis; their mean age was 37 years, mean weight of 42.1 kg and 79% were men. The second cohorts consists of 20 patients with pulmonary tuberculosis who are also HIV-positive with a mean age of 34 years; mean weight of 44 kg and 77% were men

HIV-positive patients took part in trials comparing 6 and 9 months of TB treatment. Patients were treated with isoniazid (600 mg), rifampicin (450-600 mg), pyrazinamide (1500 mg) and ethambutol (1200 mg) three times a week for the first 2 months (2HRZE_3_), followed by isoniazid and rifampicin three times a week for 4-7 months (4-7HR_3_). At the time of the study HIV-positive patients in India did not have access to antiretroviral treatment. Among HIV-negative patients one half were treated with the standard regimen (2HRZE_3_/4HR_3_), while the other half received additional ofloxacin during the intensive phase as part of a treatment shortening trial [10]. Patients were followed for three years with regular, monthly for the first 2 years and 3 monthly for the final year, sputum screening for acid-fast bacilli (AFB) and subsequent Lowenstein-Jensen (LJ) culture if found to be positive. Three sputum samples were collected at baseline and every month during treatment and 2 sputum samples were collected every month during the follow-up period.

Recurrence is defined when patients presented AFB-negative and culture negative results from sputum examination during the initial treatment followed by AFB-positive and culture positive results during follow-up sputum examination. Patients with recurrent pulmonary tuberculosis underwent a repeat of the treatment regime.

### Molecular typing and genome analysis

For the initial study DNA was extracted from *M. tuberculosis* cultures for spoligo- and MIRU-12 typing using standardised methods [9]. For the current study the banked DNA extracts, which were stored at -80 °C were measured for concentration and quality before sequencing on Illumina HiSeq WGS, ensuring a mean 100x read depth across the sequenced *M. tuberculosis* genome. FastQC v0.11.5 was used to check the quality of the reads and read trimming was performed with Trimmomatic v0.38; only reads with a Phred Score >50 were included in the single nucleotide polymorphism (SNP) analysis. Sequences were made available on the NCBI Short Read Archive (accession number **PRJNA613351**).

Separate HIV-positive (40 genomes; 20 isolate pairs) and HIV-negative (42 genomes; 21 isolate pairs) datasets were created, and phylogenetic trees built for each dataset. An alignment of high-quality SNPs was generated using Reddog (http://github.com/katholt/RedDog) with *M. tuberculosis* H37Rv as reference (Genbank accession no. **NC_000962.3**). Repetitive genetic elements were masked, and all heterozygous SNPs removed before SNP alignment was performed using Bowtie. SNP calling was preformed using BCFtools with minimum read depth of 50x. The phylogenetic tree was constructed with RAxML (GTR-GAMMA model) v8.2 using 1000 bootstrap replicates.

### Detection of drug resistance and comparison of paired isolates

Genomes were screened for markers of resistance to first-line drugs (rifampicin, isoniazid, pyrazinamide and ethambutol) using a list of high-confidence SNP mutations [11]. Snippy v.4.3.5 (http://guthub.com/tseemann/snippy) with default settings was used to detect SNPs with at least 10x read coverage that differed from the reference set with a frequency ≥0.9. Snippy allows for the detection of heterozygous SNPs and all SNPs associated with drug resistance were manually reviewed and confirmed by visual inspection of the BAM file using Artemis. A custom Perl script was employed to compare SNPs in paired isolates. Cases were classified as relapse if the isolate from the first episode differed from the second isolate by <5 SNPs. Statistical analysis was performed using the Fisher exact test to assess differences in relapse and re-infection rates.

## Results

Among HIV-negative patients (P01 to P21) with paired samples (S1 and S2), 19/21 (90%) experienced TB relapse (Figure 1A). The opposite was observed in HIV-positive patients (P22 to P41) where the vast majority of TB recurrences (16/20; 80%) resulted from reinfection (Figure 1B); the difference in frequency of reinfection was statistically significant (p=0.0001). Two paired isolates (P19-S1/S2 and P33-S1/S2) with each with 8 SNP differences were classified as re-infection using a stringent 5 SNP cut-off, despite identical spoligotype and MIRU-12 profiles (Tables 1 and 2).

**Table 1:**
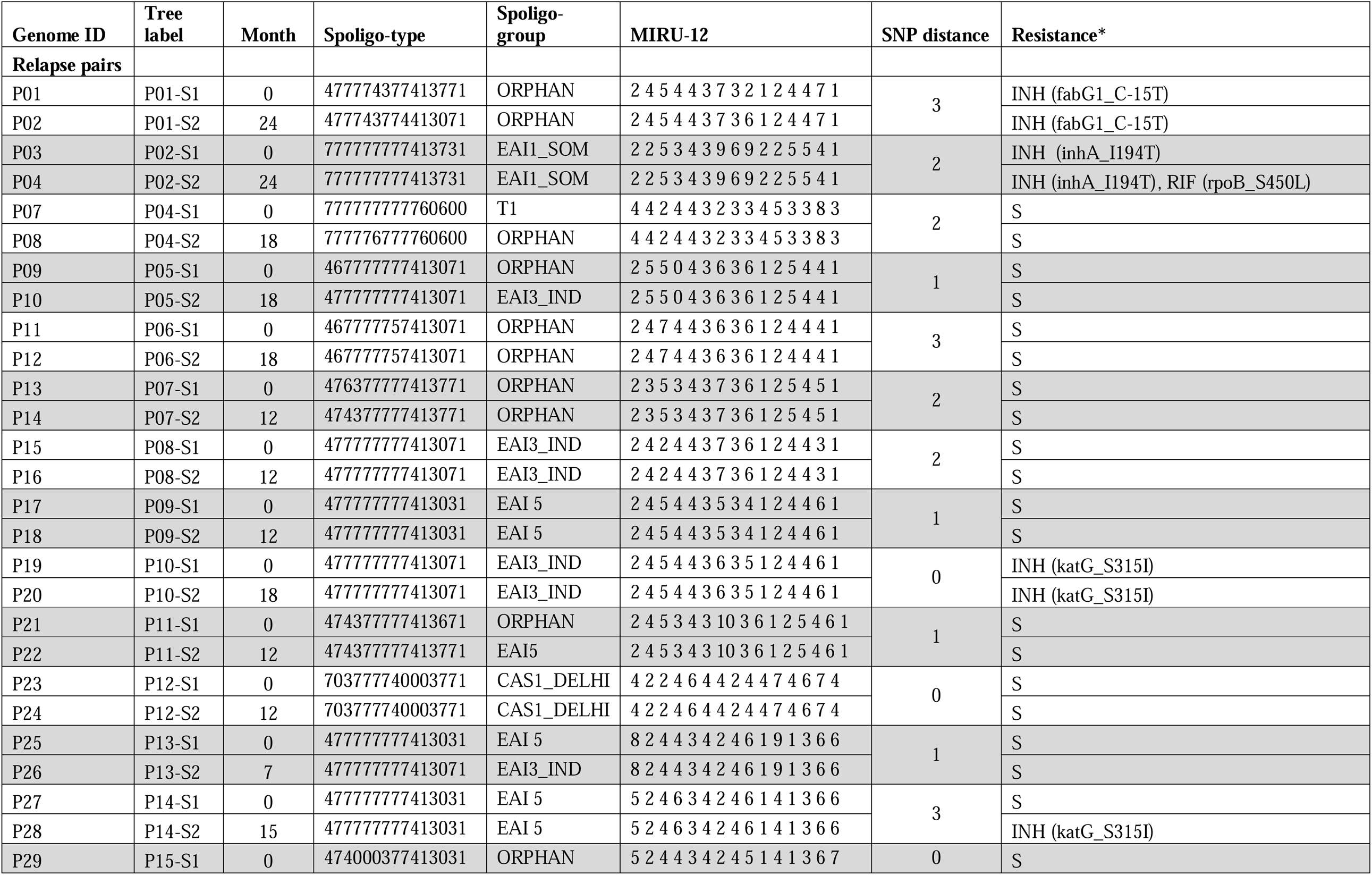
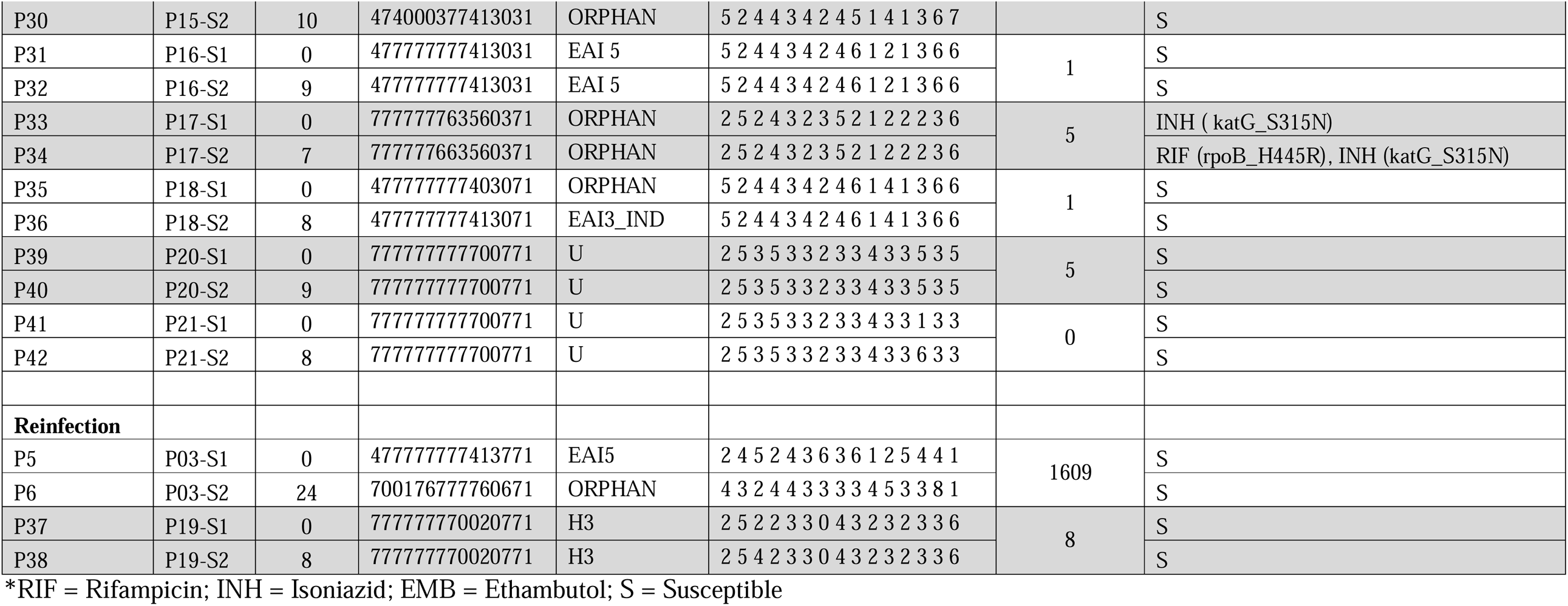
Paired *M. tuberculosis* isolates from HIV-negative patients with TB recurrence that were sequenced

**Table 2:**
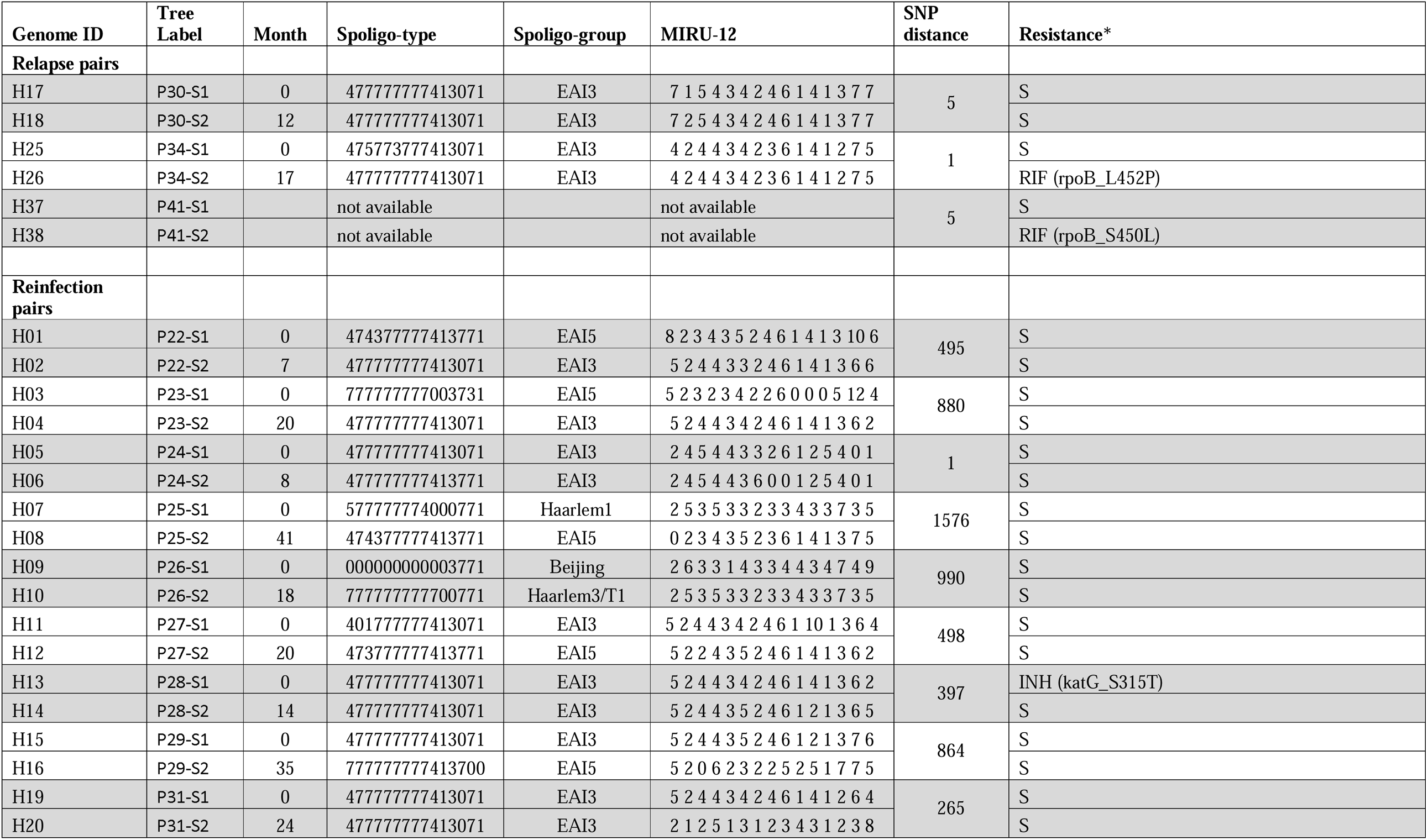
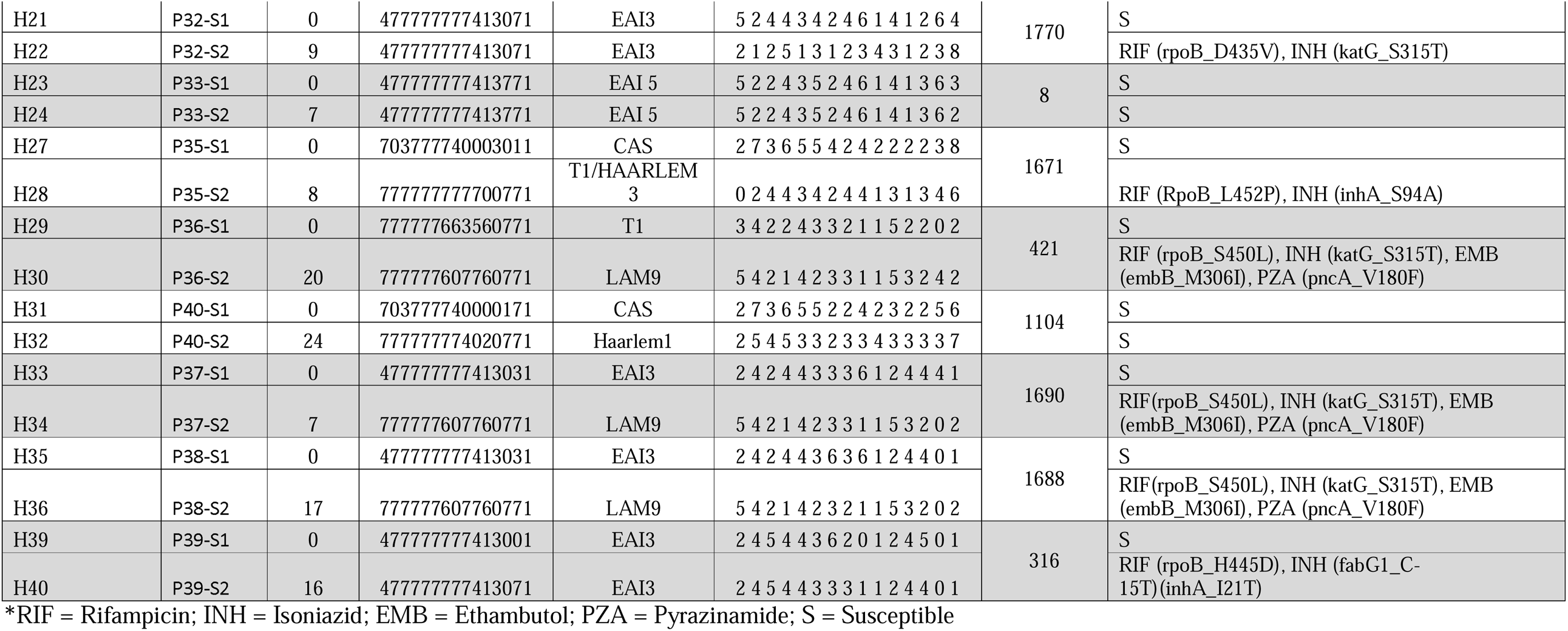
Paired *M. tuberculosis* isolates from HIV-positive patients with TB recurrence that were sequenced

**Figure 1:**
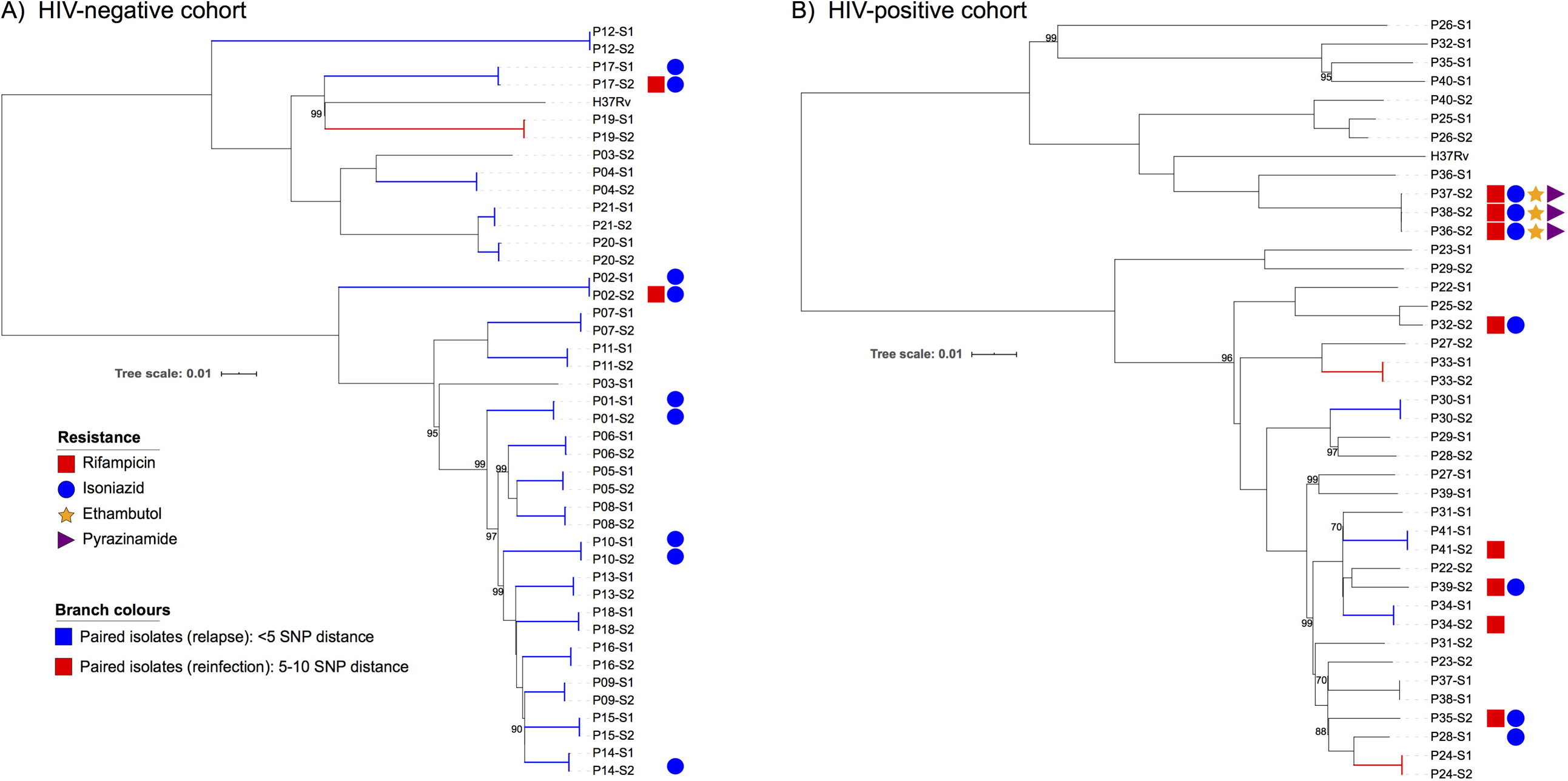
Phylogenetic tree of paired *M. tuberculosis* isolates from HIV-negative patients (panel A) and HIV-positive patients (panel B). Coloured shapes indicate resistances to antibiotics based the presences of confirmed resistance mutations in the genome sequences. Blue branches show relapse pairs >5 SNP differences. Red branches indicate re-infection pairs by strains that have 5-10 SNP difference or different MIRU-12 profile. Bootstrap values are shown on branch nodes with less than 100% support. Tree scale is the mean number of nucleotide substitutions per site.

Among HIV-negative cases reinfection was rare (2/21; P03 and P19; Figure 1A). Pre-existing mutations associated with drug resistance were documented in four relapse cases (P01, P02, P10, P17). These were *katG, inhA or fabG1* gene mutations associated with high-level isoniazid resistance (Table 1). In two instances drug resistance amplification was observed, with newly acquired rifampicin resistance conferring mutations identified in the *rpoB* gene (P02-S2; S450L and P17-S2; H445R). One isolate (P14-S2) acquired high-level isoniazid resistance via a *katG* gene mutation (S315I).

Among HIV-positive cases relapse was rare (4/20; Figure 1B). Although P37-S1 and P38-S1 were clustered together they were recovered from different patients, suggesting a transmission link or laboratory cross-contamination. Since processing of these specimens were separated in time and space, cross-contamination was considered unlikely. Likewise, isolates P36-S2, P37-S2 and P38-S2 clustered together suggesting transmission of multidrug-resistant (MDR) strain. A high percentage of re-infections 6/17 (35%) were caused by strains resistant to both isoniazid and rifampicin (Table 2). One relapse case (P24-S1/S2) had a single SNP, but was classified as a re-infection due to three differences in the MIRU-12 profile. Drug resistance amplification was demonstrated in two relapse pairs (P42-S1/P42-S2 and P34-S1/P34-S2) where the second isolate acquired a new *rpoB* mutation (S450L).

## Discussion

This is the first report to describe the added insight gained by using WGS for high-resolution differentiation of re-infection and relapse among Indian patients with TB recurrence. Previous analyses identified re-infection as the main contributor to TB recurrence in HIV-positive patients [9, 12], but WGS emphasized the fact that re-infection with MDR strains is a particular concern. Overall, WGS identified slightly increased rates of re-infection compared to those observed with MIRU-12 and spoligotyping and provided important insight into local transmission dynamics and drug resistance mutations. The high percentage of re-infection with MDR strains and the MDR-TB transmission cluster identified among HIV-positive cases, highlight the potential risk of nosocomial MDR-TB transmission linked to HIV care, as has been described in multiple settings [12].

The acquisition of rifampicin resistance in two of the three HIV-positive cases with TB relapse is also a concern and emphasizes the need to ensure optimal drug exposure, especially to rifampicin. The classification of two isolate pairs with <10 SNPs difference as re-infection, while they had identical spoligotype and MIRU-12 profiles, may be viewed as problematic. However, there is consensus that a <5 SNP cut-off is appropriate in high transmission environments, given that similar strains may be isolated following reinfection from the patient’s immediate environment. Other studies have incorporated detailed epidemiological and geographical data to resolve these ‘ambiguous’ cases [4], but similar data were not available in our study. However, the fact that paired isolates were collected 7- and 8-months apart favours re-infection, given the slow mutation rate of *M. tuberculosis*. All other re-infection pairs were unambiguous and had differences of >100 SNPs.

Our findings in HIV-negative patients are broadly concordant with previous findings that attributed most TB recurrence to relapse, using a similar 5 SNP cut-off [13]. However, few studies explored differences according to HIV infection status. A high proportion of HIV-negative relapse cases had pre-existent isoniazid mono-resistance and two of these have acquired additional rifampicin resistance at the time of relapse. Initial isoniazid resistance is well documented as the ‘common pathway’ to the emergence of multidrug resistant strains within the community [14], as is increased relapse rates among these patients [15]. While the use of Xpert MTB/RIF and other frontline genomic tests increase MDR-TB detection, their failure to diagnose isoniazid monoresistance remains a major challenge. One relapse case (P24-S1/S2) was previously classified as a re-infection given differences in both the MIRU-12 and spoligotype profiles. Strain specific differences are rarely missed by WGS, since it assesses SNPs across most of the genome. However, given that repetitive and hypervariable parts of the genome, including sections used for MIRU and spoligotyping, are routinely excluded from SNP analysis this is not completely unexpected. Consistency in the MIRU-12 and spoligotyping readouts indicates that this may represent a ‘true reinfection’ misidentified by WGS.

In conclusion, WGS based analysis confirmed previous differences observed in the relative frequency of relapse and reinfection, depending on HIV status. However, it provided enhanced insight into drug resistance acquisition and spread, with major concern related to potential MDR-TB transmission in HIV care settings.

## Acknowledgement

The authors would like to acknowledge Danielle Somers and the Office of Global Health at the University of Sydney for assisting with the work preformed in this study.

